# Neural activity is modulated by spontaneous and volitionally controlled breathing

**DOI:** 10.1101/2023.08.30.555484

**Authors:** Suvi Karjalainen, Jan Kujala, Tiina Parviainen

## Abstract

Recent studies have provided evidence regarding respiration-brain coupling, but how continuously varying dynamics of breathing modulate neural activity is not known. We examined whether the neural state differs between spontaneous and volitionally controlled breathing and across the phases of breathing, inspiration and expiration. Magnetoencephalography with a respiratory belt was used to record cortical oscillatory activity during spontaneous, deep, and square breathing (n=33). Alpha power was suppressed during inspiration and increased during expiration (p<0.01) indicating dynamically fluctuating neural states across the respiratory cycle. Compared to spontaneous and square breathing, alpha power increased during deep breathing (p<0.01). We also observed a steeper aperiodic slope and a broadband shift in the power spectrum during square breathing in comparison with spontaneous breathing suggesting that spectral characteristics of neural activity are modulated by the rate, depth, and pattern of breathing. Altogether, we demonstrate that neural activity is modulated by breathing techniques and phases of breathing.

## Introduction

Breathing is a vital, cyclic bodily rhythm consisting of active inspiration and passive expiration. Although autonomic breathing is mostly driven by metabolic demands, breathing can also be volitionally controlled to adapt to changes in internal motivational states, such as emotions, or in external environmental demands (Homma & Masaoka, 2008). Practices that utilize volitional regulation of breathing (e.g., rate, depth, and pattern of breathing, inspiration/expiration ratio) have been shown to induce behavioral, psychological, autonomic, and neural changes (Lin, Tai, & Fan, 2014; Song & Lehrer, 2003; Zaccaro et al., 2018). Although being used as a therapeutic technique for thousands of years, the interaction between respiration and central nervous system activity during the implementation of different breathing techniques, such as deep and square breathing, is not well understood. Given that such neural modulation by breathing exists, it would entail volitional modulation of neural activity by controlled breathing. Understanding of this modulation could yield a mechanistic account for the behavioral and clinical implications for which there is some initial evidence, such as facilitation of optimal performance and learning or improved treatment of psychiatric symptoms.

Key neural elements for generating breathing patterns involve ponto-medullary network structures, and especially the role of a medullary microcircuit, preBötzinger complex, is fundamental in coordinating the phases of the breathing cycle (Del Negro, Funk, & Feldman, 2018; Yang & Feldman, 2018). However, in volitional control of breathing, more extended network of subcortical and cortical structures is involved (Guz, 1997; McKay, Evans, Frackowiak, & Corfield, 2003). For instance, voluntary pacing of breathing rate has been shown to engage frontotemporal-insular network, whereas awareness of breathing seems to be associated with the activation of anterior cingulate, insular, hippocampal, and premotor cortices (Herrero, Khuvis, Yeagle, Cerf, & Mehta, 2018).

Substantial evidence in animal studies, as well as emerging findings also concerning humans, show that the electrophysiological signaling in the brain is governed by the respiratory signal (Karalis & Sirota, 2022; Ito et al., 2014). Specifically, nasal respiration has been shown to entrain neural oscillations at the same frequency as breathing in several regions of the rodent brain (Corcoran, Pezzulo, & Hohwy, 2018; Tort, Brankačk, & Draguhn, 2018). Intracranial electroencephalography (EEG) recordings in humans have revealed that natural breathing synchronizes neural activity in the piriform cortex and limbic-related brain regions, such as hippocampus and amygdala, especially during nasal breathing, thus highlighting the importance of the nasal route in generating respiration-related oscillatory activity (Zelano et al., 2016). Moreover, respiration-modulated neural oscillations have been observed across several frequency bands within a network consisting of both cortical and subcortical brain areas (Kluger & Gross, 2021). Besides spontaneous breathing, cortical oscillatory activity seems to be influenced by volitional control of breathing (e.g., Kluger & Gross, 2020; Schellart & Reits, 1999). Based on these findings, it has been suggested that respiration-brain coupling may be a mechanism facilitating both interregional communication and local information processing in the human brain (Heck et al., 2017; Tort et al., 2018). Thus, respiration-brain coupling has been regarded as a hierarchical principle for organizing neural oscillatory activity throughout the human brain (Herrero et al., 2018).

Within the cyclic rhythm of breathing the phases of breathing (i.e., inspiration and expiration) have been found to modulate the dynamics of neural oscillatory activity (Bušek & Kemlink, 2005; Hsu, Tseng, Hsieh, & Hsieh, 2020). On the other hand, opposing findings showing that cortical activity is not modulated across the respiratory phases have also been reported (e.g., Park, Barnoud, Trang, Kannape, Schaller, & Blanke, 2020). While the findings regarding modulatory effects of respiratory phases on neural activity remain scarce and partly contradictory, there is compelling evidence indicating that sensory, motor, cognitive, and emotional processes are modulated across the respiratory cycle. For instance, the magnitude of startle eye-blink (Schulz, Schilling, Vögele, Larra, & Schächinger, 2016), detection of visual signals (Flexman, Demaree, & Simpson, 1974), reaction times (Johannknecht & Kayser, 2022), initiation of voluntary actions (Park, Barnoud, Trang, Kannape, Schaller, & Blanke, 2020) as well as fear discrimination and memory retrieval (Zelano et al., 2016) have been observed to vary with the phases of respiration. Based on these findings, it seems fair to say that phases of respiration influence a variety of processes ranging from basic reflexes to higher information processing. However, the question remains whether the modulation of the aforementioned processes is paralleled by modulation of neural activity reflecting different neural states during inspiration and expiration.

Some evidence exists that breathing-induced modulations in neural synchrony underlie the modulatory effects observed in perceptual and cognitive functions (Varga & Heck, 2017). Initial findings regarding respiration-brain-cognition coupling have shown that respiration modulates perceptual sensitivity through modulations in posterior alpha power that is regarded as a marker of neural excitability (Kluger, Balestrieri, Busch, & Gross, 2021). Similarly, spontaneous inspiration at the onset of a cognitive task has been observed to be accompanied by a shift in functional neural network architecture likely facilitating improved performance in a visuospatial task (Perl et al., 2019). There is also initial empirical evidence suggesting that there are ‘optimal’ states within different bodily rhythms (diastole for cardiac cycle and expiration for respiratory cycle) that facilitate learning-related processes at the neural level (Waselius, Wikgren, Penttonen, & Nokia, 2019; Waselius, Xu, Sparre, Penttonen, & Nokia, 2022). According to these findings, neural states seem to dynamically fluctuate depending on the phase of the respiratory cycle and these fluctuations seem to be also behaviorally relevant.

In this study, we investigated how breathing modulates the functional state of the brain. So far, the emerging evidence has demonstrated modulatory effects of breathing on neural activity and behavior, but the literature regarding non-invasive neuroimaging studies in humans is still very scarce. Especially, whether volitional control of breathing rate, depth, and pattern can change the state of brain signaling has received little attention so far. Furthermore, there is a lack of studies investigating whether the phases of breathing modulate neural activity in the same manner during both spontaneous and volitionally controlled breathing. Thus, we examined how neural activity differs between spontaneous and volitionally controlled deep and square breathing. We also investigated whether and how neural activity is modulated across the phases of breathing. Specifically, we examined whether the transitions from inspiration to expiration and from expiration to inspiration during both spontaneous and volitionally controlled breathing coincide with a shift in the state of the brain.

Considering neural oscillations as functional markers of brain states (Buzsáki & Draguhn, 2004; Thut, Miniussi, & Gross, 2012) we recorded continuous electrophysiological cortical activity using magnetoencephalography (MEG) together with a respiratory belt from 40 participants during three breathing conditions: 1) spontaneous, 2) deep, and 3) square breathing. We especially focused on the breathing-related modulations in alpha-band oscillatory activity (∼ 10 Hz), one of the most prominent rhythms in the human brain. This choice was motivated by the observations that alpha-band oscillations play an active role in cognitive processes, such as functional inhibition and allocation of attentional resources (Foxe & Snyder, 2011; Jensen & Mazaheri, 2010; Klimesch, 2012; Klimesch, Sauseng, & Hanslmayr, 2007). Thus, using alpha-band oscillatory activity as a proxy of the neural states, we assessed how different breathing techniques as well as phases of breathing modulate the functional state of the brain. Since neural activity consists of not only periodic, oscillatory components but also aperiodic components typically showing 1/f-like characteristics (Donoghue et al., 2020; He, 2014), we also disentangled the differences in aperiodic activity between breathing techniques.

## Results

### Respiration

To assess how breathing differed across breathing techniques, the respiratory rate was calculated for each technique: spontaneous, deep, and square breathing. Respiratory rate varied across breathing techniques as follows: spontaneous 13.34 ± 3.84, deep 5.30 ± 2.19 and square breathing 8.03 ± 3.87 (breaths/min, mean ± SD). There was a statistically significant difference in respiratory rate between breathing techniques χ2(2) = 49.636, p < 0.001. Post hoc analyses indicated that spontaneous breathing differed from volitionally controlled deep (Z = −5.012, p < 0.001) and square breathing techniques (Z = −4.476, p < 0.001). Also, the volitionally controlled deep and square breathing techniques differed from each other (Z = −3.976, p < 0.001). Fig. 1 illustrates that the respiratory rate was higher during spontaneous breathing in comparison with volitionally controlled deep and square breathing techniques. Moreover, respiratory rate was higher during square than during deep breathing. These findings confirm that the participants were able to perform the breathing tasks as instructed.

**Figure 1.**
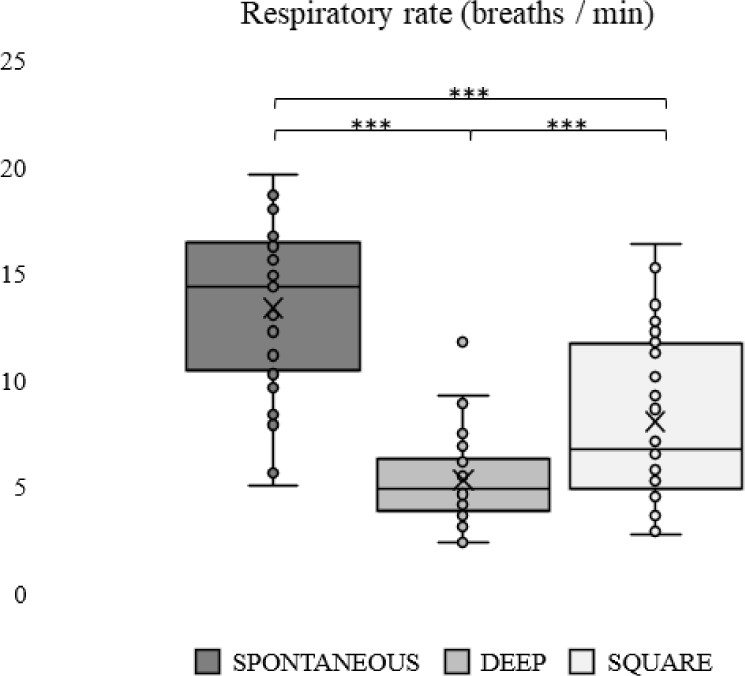
Mean respiratory rate during the implementation of each breathing technique. In the box plots the upper and lower box boundaries represent the 75^th^ and 25^th^ percentiles, respectively, whiskers denote minimum and maximum values, cross the mean value, line inside the box the median value, and circles outside the whiskers are considered outliers. *** p < 0.001

### Periodic and aperiodic activity across breathing techniques

To investigate whether and how the state of the brain electrophysiological activity differs between breathing techniques, we investigated the spatial distribution of modulations in periodic and aperiodic neural activity during spontaneous breathing and the two techniques of volitionally controlled breathing. Fig. 2 depicts the spatial distribution of statistically significant differences in periodic (alpha power, upper row) and aperiodic (exponent, middle row; offset, bottom row) activity between breathing techniques. The parcel-level dependent samples t-test results (p < 0.01, corrected for multiple comparisons) demonstrated that alpha power was higher during deep breathing in comparison with spontaneous and square breathing. When deep and spontaneous breathing were compared, these differences in alpha power were observed bilaterally, especially in the occipital and temporal areas, but extending also to parietal and prefrontal areas as well as cuneus and cingulate cortex. When comparing deep and square breathing, increase in alpha power was limited to the left temporal and inferior frontal areas, with a cluster also in the left superior frontal gyrus. No statistically significant differences in alpha power were found between spontaneous and square breathing. The findings regarding aperiodic activity showed that the exponent was higher and offset lower during square breathing in comparison with spontaneous breathing in the left superior frontal gyrus. No other statistically significant differences in aperiodic activity between breathing techniques were found.

**Figure 2.**
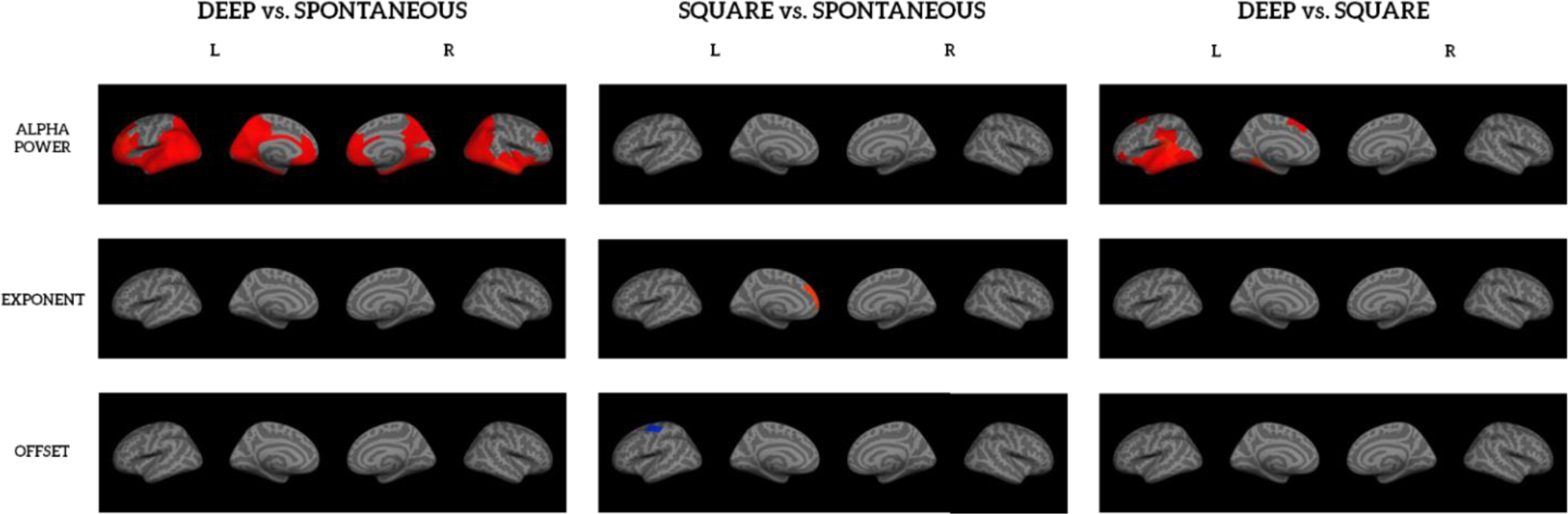
Modulations of periodic and aperiodic activity between different breathing techniques. Statistically significant differences (p < 0.01, corrected for multiple comparisons) were observed in periodic and aperiodic activity between breathing techniques. Blue color indicates statistically significant decreases and red color statistically significant increases in periodic and aperiodic activity between breathing techniques.

Since breathing-related head movements can a be potential confound in our analysis, we examined whether the differences in periodic and aperiodic activity between breathing techniques could be explained by differences in head movements. We used the head position information collected continuously during the MEG recordings (n = 31, continuous head position information not available for two participants) to calculate the mean movement velocity (sum of vectors in x, y, and z directions) across timepoints. Thereafter, we investigated whether the difference in mean movement velocity (mm/s) between breathing techniques was correlated (Spearman’s Rank correlation) with the difference in neural activity within the parcels in which statistically significant differences between breathing techniques were observed. The findings suggest that only in two parcels in the left anterior temporal lobe the difference in alpha power between deep and spontaneous breathing could be explained by artefacts or variance in the signal caused by breathing-related head movements (p < 0.05, uncorrected). No other correlations were found. Thus, the results regarding left anterior temporal areas need to be interpreted with caution.

### Modulation of alpha power across the phases of breathing

To explore whether neural activity is modulated across the phases of breathing the spatial distribution of suppressions and increases of alpha-band oscillatory activity during inspiration and expiration were investigated. Results of the source reconstruction of oscillatory power modulations across the phases of respiration are illustrated in the Fig. 3. Overall, the source-level cluster-based permutation test results (p < 0.01, corrected for multiple comparisons) indicated that alpha power decreased during inspiration and increased during expiration in the time windows from 0 to 2000 ms with respect to the onsets of inspiration and expiration, respectively, in comparison with the baseline time window of - 500 to 0 ms during spontaneous and deep breathing.

**Figure 3.**
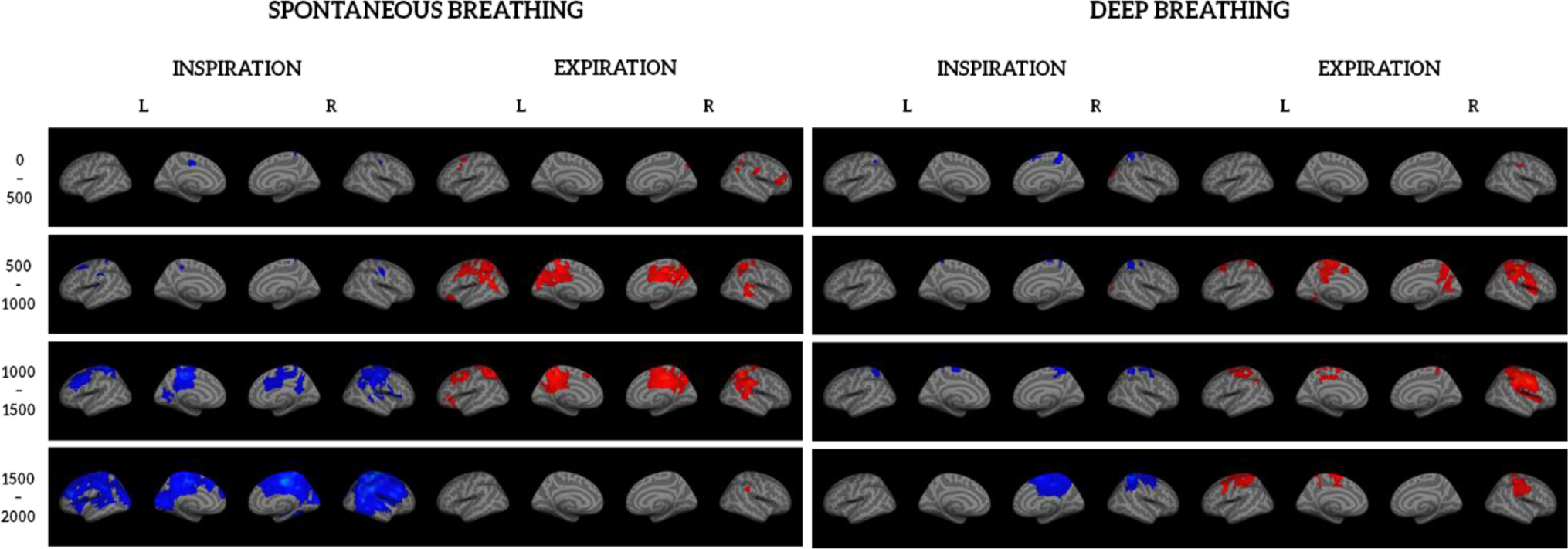
Alpha power modulations across the phases of respiration. Source-level cluster-based permutation test results indicated that alpha power (8 – 13 Hz) decreased during inspiration and increased during expiration in the time windows from 0 to 2000 ms with respect to the onset of inspiration and expiration in comparison with the baseline time window of −500 to 0 ms especially in the sensorimotor and parietal areas during spontaneous and deep breathing (p < 0.01, corrected for multiple comparisons). Blue color indicates statistically significant decreases and red color statistically significant increases in the alpha power in comparison with the baseline time window.

During spontaneous breathing at the early stage of inspiration (time window 0 – 1500 ms), alpha power decreased bilaterally in the sensorimotor and frontal areas as well as in the precuneus and cingulate cortex. In the later stage of inspiration (time window 1500 – 2000 ms) decreases were more widespread covering most of the cortical and subcortical areas. During the earliest stage of expiration (time window 0 – 500 ms), increased alpha power was observed bilaterally in the frontal areas and in the right parietal area. During the mid expiration (time window 500 – 1500 ms) increased alpha power was observed bilaterally in the sensorimotor, parietal, and superior temporal areas. Moreover, increases were found bilaterally in the cingulate cortex, cuneus, and precuneus as well as in the left orbitofrontal cortex. During the latest stage of expiration (time window 1500 – 2000 ms) increased alpha power was observed only in the right parietal area.

During deep breathing decreased alpha power was observed bilaterally in the sensorimotor and parietal areas in the early stage of inspiration (time window 0 – 1500 ms). During the later stage of inspiration (time window 1500 – 2000 ms) decreased alpha power was only found in the right sensorimotor and parietal areas as well as right precuneus and cingulate cortex. Moreover, increased alpha power was observed during the early stage of inspiration (time windows 0 – 1000 ms) in the right occipital areas. During the earliest stage of expiration (time window 0 – 500 ms) increased alpha power was observed in the right parietal area. During the later stage of expiration (time window 500 – 2000 ms) increases in alpha power were found bilaterally in the sensorimotor and parietal areas as well as in the cuneus and precuneus. Moreover, increases in alpha power over left the occipital areas were observed in the time window from 500 ms to 1000 ms of expiration.

To summarize, during the inspiratory phase of spontaneous breathing alpha power suppression originates from the sensorimotor areas and then extends also to the fronto-temporal areas, whereas during the expiratory phase increased alpha power seems to remain mainly in the sensorimotor areas. During deep breathing alpha power modulations across the phases of breathing are limited to the sensorimotor areas.

## Discussion

The aim of this study was to explore whether the functional state of the brain dynamically fluctuates across breathing techniques and phases of breathing. We observed a robust shift in alpha power across the phases of breathing: the level of alpha power was suppressed during inspiration and increased during expiration. These effects were observed especially in the sensorimotor areas during both spontaneous and deep breathing. This finding confirms our hypothesis that the transitions from inspiration to expiration and from expiration to inspiration during both spontaneous and volitionally controlled deep breathing coincide with a shift in the state of the brain. In accordance with the present results, recent studies have demonstrated similar modulations of alpha power across the phases of breathing both in rest (Kluger & Gross, 2021) and task-based conditions (Kluger et al., 2021; Perl et al., 2019) during spontaneous breathing. Our findings provide thus confirmation for the recently emerged evidence on the impact of the dynamically changing body physiology, namely respiratory cycle, on the dynamic variation in the neural state. We extend these findings to cover also volitionally controlled deep breathing, where the alpha power levels varied essentially in the same manner as compared with spontaneous breathing across the phases of breathing.

Our findings regarding breathing techniques indicated that the power of alpha-band oscillatory activity is increased during volitionally controlled deep breathing in comparison with spontaneous and square breathing. This observation confirms and extends the initial evidence from early 2000s by Bušek and Kemlink (2005) showing that alpha power is modulated depending on the breathing frequency. Namely, they found that alpha power was significantly higher during slow breathing (0.1 Hz) in comparison with fast breathing (0.5 Hz) as well as during normal breathing (0.25 Hz) compared with fast breathing paced by a metronome (Bušek & Kemlink, 2005). Since the respiratory rate was not paced by any external stimuli in our study, the individual variability in respiratory rate especially during spontaneous and square breathing was rather high. Thus, the variability in the respiratory rate could explain why, against our expectations, no differences in alpha power were observed between spontaneous and square breathing. Although both deep and square breathing techniques aim at promoting recovery through the activation of parasympathetic division of autonomic nervous system (Zaccaro et al., 2018), only deep breathing seemed to have a modulatory effect on the oscillatory activity of the brain. Since only a few participants were familiar with the square breathing technique in comparison with deep breathing that was familiar to everyone, the implementation of square breathing may have required more attentional resources resulting in alpha power that resembles more closely the neural activity observed during spontaneous breathing than deep breathing. It is also worth noting that, based on the findings by Bušek and Kemlink (2005) it was not possible to point out exactly in which cortical areas alpha power was modulated, because the modulations in the EEG spectral power were examined at a rather coarse level across several sensors. Our approach, in turn, allowed us to investigate the respiration-related modulations in neural activity across the whole cortex in detail.

Furthermore, our results evidence that not only oscillatory activity but also the more general spectral characteristics of ongoing neural activity are modulated by breathing. The larger exponent of aperiodic activity observed during square breathing in comparison with spontaneous breathing might indicate stronger inhibitory activity as demonstrated by the steeper 1/f slope in the power spectrum (Donoghue et al., 2020; Gao, Peterson, & Voytek, 2017; He, 2014). Aperiodic offset, in turn, has been suggested to reflect a broadband shift in the power spectrum across frequencies and has been reported to be positively correlated with neuronal firing rates (Donoghue et al., 2020; Manning, Jacobs, Fried, & Kahana, 2009). Thus, lower offset observed during square breathing in comparison with spontaneous breathing could reflect higher levels of neuronal firing rates during spontaneous breathing.

Interestingly, these effects were only observed in spatially limited areas in the left superior frontal gyrus, suggesting that volitionally controlled square breathing could modulate inhibitory activity and neuronal firing rates locally in the frontal brain areas compared to spontaneous breathing. Since superior frontal gyrus has been observed to be anatomically and functionally connected with nodes of the default mode, cognitive control, and executive networks (Li et al., 2013) and as it has also been shown to play a role in cognitive processes such as working memory, planning, and problem solving as well as motivation (Szczepanski & Knight, 2014), it would be intriguing to explore whether breathing-related modulations in neural activity would also be associated with these aforementioned processes. All in all, the findings regarding periodic and aperiodic activity support the assumption that the functional state of the brain is modulated by breathing. Yet further research is required to deepen our understanding of the dynamic interactions between the spectral features of neural oscillatory activity during spontaneous and volitionally controlled breathing.

Altogether, our findings clearly demonstrate that the functional state of the brain is modulated by the phases of breathing and can also be volitionally influenced by varying the rate, depth, and pattern of breathing. The regular pattern of dynamic fluctuations in alpha power across the phases of breathing covered mainly sensorimotor areas representing a general cycle-by-cycle effect irrespective of the breathing technique. Interestingly, deep breathing technique seemed to additionally modulate neural activity within a cortical network extending to frontal and temporal areas.

The spatial distribution of breathing-related modulations in alpha power observed in our study seems to overlap with the sources of respiration-modulated brain oscillations identified by Kluger and Gross (2021), including superior temporal gyrus, temporal pole, and cingulate cortex. Our findings are also consistent with the very few studies that have investigated the modulatory effects of volitionally controlled breathing on neural activity (Herrero et al., 2018; McKay et al., 2003) showing that volitional control of breathing engages frontotemporal and sensorimotor cortices. Thus, the results of this study add to the existing evidence indicating that besides brainstem structures (Del Negro, Funk & Feldman, 2018; Yang & Feldman, 2018), also cortical areas are involved in the act of breathing. Since continuous streams of exteroceptive (e.g., auditory, mechanical, and thermal sensations of air flow through the nose), proprioceptive (e.g., chest movements), and interoceptive (e.g., CO2 and O2 level monitoring) sensory input are provided to the brain during breathing, the widespread breathing-related modulations of neural activity may reflect the complex interaction between these parallel perceptual processes. It is not, however, feasible to tell apart whether these modulations are associated with the perception of breathing-related sensory input or the sensorimotor control of breathing. Hence, the question of to what extent each of these processes are involved in spontaneous and volitionally controlled breathing remains to be explored in greater detail in the future.

Strong evidence provided by previous studies suggests an inverse relationship between alpha power and neural excitability (Chapeton, Haque, Wittig, Inati, & Zaghloul, 2019; Hanslmayr, Gross, Klimesch, & Shapiro, 2011; Iemi et al., 2022; Kluger et al., 2021). Our results would thus suggest lower cortical excitability during expiratory phase compared to inspiratory phase of breathing, as well as during deep breathing as compared with spontaneous and square breathing.

Another interpretation suggests that modulations of alpha power reflect fluctuations between externally and internally oriented neural states in which brain is biased either towards processing information from the external or internal world (see review Hanslmayr et al., 2011). To be more specific, decreases in alpha amplitude have been regarded as externally oriented states, whereas increases in alpha amplitude have been considered to indicate internally oriented states (Hanslmayr et al., 2011; Knyazev, Slobodskoj-Plusnin, Bocharov, & Pylkova, 2011). Thus, the alpha power decrease during inspiration and increase during expiration observed in our study would implicate externally and internally oriented states, respectively. Similarly, higher alpha power during deep breathing in comparison with spontaneous and square breathing could reflect a more internally oriented state. This interpretation is consistent with recent evidence demonstrating that the amplitude of heart evoked potentials (i.e., a marker of interoceptive information processing) is higher during expiration in comparison with inspiration suggesting a shift towards internally oriented state facilitated by expiration (Zaccaro, Perrucci, Parrotta, Costantini, & Ferri, 2022).

These two interpretations regarding cortical excitability and internally vs. externally oriented states may be considered complementary to each other. The idea of internally and externally oriented states is supported by the observations that alpha-band oscillatory activity, especially a task-related decrease in alpha power, can predict whether an external stimulus is perceived or not (e.g., Hanslmayr et al., 2011; Kluger et al., 2021; Perl et al., 2019). On the contrary, increase in alpha power has been frequently linked with internal mental processes, such as self-referential thoughts, imagination, mental imagery, and internal attention (e.g., Cooper et al., 2003; Klinger et al., 1973; Knyazev et al., 2011; Ray and Cole, 1985). The enhanced alpha activity observed during internally oriented information processing has also been shown to be disrupted when shifting to the processing of external events (Knyazev et al., 2011). This observation further highlights the relevance of alpha-band oscillatory activity in the fluctuation between internally and externally oriented states. Combined with the robust evidence indicating an inverse relationship between alpha power and neural excitability (e.g., Chapeton et al., 2019; Iemi et al., 2022), these findings together suggest that higher alpha power is accompanied by lower cortical excitability and internally oriented information processing, whereas lower alpha power is associated with higher cortical excitability and externally oriented information processing.

Although the understanding of how breathing modulates the state of the brain is constantly advancing, the intriguing question of why such coupling between respiration and brain activity exists remains unanswered. Physiologically, one possible explanation relies on cardiorespiratory coupling, the universally observed interaction between respiratory and cardiac activity, which is suggested to be fundamental in facilitating the efficiency of gas exchange (Grossman & Taylor, 2007; Yasuma & Hayano, 2004). Respiration-brain coupling may further represent the dependence of brain electrophysiology on this vital metabolic coupling. From the evolutionary perspective, it has been proposed that via this metabolic coupling bodily rhythms may have been fundamental in the functional organization of the nervous system (Parviainen, Lyyra, & Nokia, 2022). As a consequence, besides efficient homeostatic regulation, respiration may also optimize the bodily and neural states to facilitate information processing. Intriguingly, it has been further speculated that the evolutionarily preserved associations between bodily and neural rhythms as well as cognitive functions within an individual may have enabled a coherent perception of the world also across individuals possibly facilitating social interaction, co-operation, and decision making (Parviainen et al., 2022). Within this framework, the ability to volitionally control the state of the body and brain by changing the rate, depth, or pattern of breathing could be meaningful in modulating the way we interact with others and the environment around us. While being still rather speculative, this view emphasizes the relevance of respiration-brain coupling also for behavior.

While this study successfully demonstrated how neural activity is modulated by spontaneous and volitionally controlled breathing, it has certain limitations. First, the relatively high inter-individual variability in the respiratory rate, especially during spontaneous and volitionally controlled square breathing, may have affected our findings. However, to enhance the practical and clinical relevance of this study, it was crucial to use natural, self-paced breathing instead of pacing the breathing rate and duration of respiratory phases using a metronome, for instance. Besides the breathing rate, depth and pattern of breathing were also volitionally controlled during deep and square breathing in the current study. Therefore, it is likely that these factors contributed to the modulation of neural activity during different breathing techniques as well.

Although the sample size in our study was comparable with other studies in the field (e.g., Kluger & Gross, 2021, Kluger et al., 2021), larger samples in the future would be beneficial in establishing the inter- and intra-individual variability of respiratory-brain coupling. With regard to the research methods, limitations related to source reconstruction need to be acknowledged. Due to the limited spatial resolution of MEG and challenges posed by localization of deeper sources (Gross, et al., 2013; Hari, Joutsiniemi, & Sarvas, 1988), it was not feasible to examine the modulatory effects of breathing in subcortical structures, such as insula, limbic system, or cerebellum, in detail. Additional uncertainty arises from the use of a template in the source-level analyses of MEG data, because individual anatomical magnetic resonance images (MRI) were not available.

Since breathing techniques are widely used in therapeutic settings, meditation, self-regulation, and optimizing performance (e.g., police officers, military personnel, athletes), this study improves our understanding of the beneficial effects of volitionally controlled breathing techniques on the central nervous system level. These fundamental findings also open an important avenue of studies to better understand the neural effects of atypical breathing patterns and disrupted breathing observed in various neurodevelopmental (e.g., autism spectrum disorder) and psychiatric conditions (e.g., panic disorder) (Ming, Patel, Kang, Chokroverty, & Julu, 2016; Nardi, Freire, & Zin, 2009). Future studies with detailed measures of breathing (e.g., oxygen and carbon dioxide, tidal volume) are warranted to develop a broader picture of the coupling between breathing and neural activity. Furthermore, a greater focus on different types of breathing (e.g., nasal vs. oral, breath holding, hyperventilation) during both rest and task contexts could provide valuable knowledge on to what extent breathing modulates the functional state of the brain.

## Conclusions

This work contributes to the rapidly expanding knowledge of respiration-brain coupling by evidencing how spontaneous and volitionally controlled breathing modulate the functional state of the brain. We showed a robust shift from suppressed alpha power during inspiration to increased alpha power during expiration. Compared to spontaneous and square breathing, an increase in alpha power during deep breathing was found. We also observed higher exponent and lower offset during square breathing in comparison with spontaneous breathing. Altogether, these findings indicate that the spectral characteristics of ongoing neural activity are modulated by breathing techniques and phases of breathing. Further research is needed to establish the possible behavioural and clinical implications of the interaction between breathing and neural activity.

## Acknowledgements

S.K. discloses support for the research of this work from the Alfred Kordelin Foundation, the Finnish Cultural Foundation, and the Central Finland Regional Fund of the Finnish Cultural Foundation. The funders had no role in study design, data collection and analysis, decision to publish, or preparation of the manuscript. The authors wish to express their thanks to Viki-Veikko Elomaa for developing the strain gauge breathing sensor and Pessi Lyyra for providing us helpful advice in the early stages of this project. We are also grateful to Hanna Honkanen, Hanna-Maija Lapinkero, Sannamari Matveinen, Emilia Penttinen, and Reetta Siikavirta for their assistance during data collection.

## Materials and methods

### Participants

Data used in this study consists of two data sets acquired in 2018-2019 (n = 12) and 2020-2021 (n = 31). A total of 43 participants were recruited via mailing lists and posters. Due to Covid-19 pandemic, some visits to the MEG laboratory had to be rescheduled or cancelled resulting in a drop out of three participants. Only participants with a complete MEG data set of adequate quality were included in the analyses and thus, the final sample consisted of 33 participants (24 female, age 25.39 ± 6.87 years [mean ± SD], range 19-57 years). All participants were screened for the following exclusion criteria: history of cardiovascular or respiratory disease, head trauma, intellectual disability, neurological or psychiatric condition, use of medication affecting the nervous system, and claustrophobia. Moreover, participants included in this study were 18-65 years old and had normal or corrected-to-normal hearing and vision. Written informed consent was obtained from all participants and a gift card (15 €) was given as a reward for taking part in the study. Ethical approval was obtained from the local Ethics Committee of the University of Jyväskylä and the study was conducted in accordance with the Declaration of Helsinki.

### Procedure

Since the first set of data collection did not include cognitive tests, only one visit in the MEG laboratory of Centre for Interdisciplinary Brain Research (Jyväskylä, Finland) was required (see details of the MEG measurement below). During the visit participants were informed about the study procedures and an informed consent form was signed. Thereafter recordings of central and autonomic nervous system activity were conducted, and questionnaires were given to the participants to be filled out at home. The second set of data collection consisted of two visits to the Centre for Interdisciplinary Brain Research. During the first visit (1.5 hours) participants were informed about the study procedures and an informed consent form was signed. Following these preparations, cognitive tests were conducted, and a set of questionnaires were given to the participant to be filled out at home in between the laboratory visits. The second visit (2.5 hours) included recordings of central and autonomic nervous system activity. The participants were instructed to refrain from caffeine and nicotine for 3 hours prior to the visit. MEG data collection was conducted either in the morning (9-12 am) or in the afternoon (12-15 pm) to minimize the effect of circadian rhythm. During the MEG recordings participants were seated upright in the MEG device (68° gantry position). To minimize head movements, especially related to breathing, participant’s position within the MEG device were stabilized with pillows and they were instructed to avoid any excessive movements or muscle contractions.

MEG recordings consisted of three breathing conditions: spontaneous breathing (8 min), deep breathing (8 min), and square breathing (8 min). In all conditions participants were instructed to sit relaxed and breathe through their nose with a respiratory rate that felt natural to them. Additionally, participants were instructed to keep their eyes on a fixation cross centered on a wall in front of them (distance from the participant’s eyes ∼ 2,5 m). For spontaneous breathing condition no specific instructions related to breathing were given. During deep breathing participants were to calmly inhale and fully fill their lungs with air and then exhale and push all the air out of the lungs. During square breathing participants were instructed to pace their breathing pattern so that all the phases including breath holding after each inspiration and expiration phase (inhale – hold – exhale – hold) were of similar duration. They were also told not to count or otherwise externally pace the phases of breathing to avoid any cognitive or motor influences. Prior to the recording of each breathing condition, the breathing technique in question was practiced for 30-60 seconds to confirm correct implementation of the technique as well as to assess the quality of the respiratory signal. Short self-paced breaks were kept between each breathing condition. To avoid any effect regarding the order of the breathing techniques, the order of deep and square breathing techniques was altered for every other participant. Experimental design also included a resting state condition with eyes closed, a heartbeat discrimination task as well as self-reports of felt arousal and feelings that are not reported here.

### Data recordings

MEG data were recorded using a whole-head 306-sensor (102 magnetometer channels and 204 planar gradiometer channels) Elekta Neuromag TRIUX system (MEGIN Oy, Helsinki, Finland) with a sampling rate of 1000 Hz and a band-pass filter of 0.1-330 Hz. MEG recordings were conducted in a magnetically shielded sound-attenuated room. During the MEG recordings the head position with respect to the MEG sensors was continuously monitored with five head position indicator (HPI) coils attached to the scalp of the participant (three HPI coils placed on the forehead and one behind each ear). For this purpose and to allow co-registration with the MRI template, the anatomical landmarks (nasion, left and right preauricular points), positions of the HPI coils, and an additional set of points (∼ 150) randomly distributed over the scalp were digitized using the Polhemus Isotrak digital tracker system (Polhemus, Colchester, VT, USA) prior to the MEG recordings.

Simultaneously with the MEG data electrocardiogram (ECG), electro-oculogram (EOG) and respiratory signals were recorded. EOG was recorded with an electrode pair placed diagonally above the right eye and below the left eye to detect eye blinks and saccades. For ECG recording one electrode was attached below each collar bone. Additionally, one ground electrode was attached to the right collarbone. The rate and phases of breathing were measured with a respiratory belt placed around the participant’s lower chest. For the first 12 participants a piezo-based breathing sensor (Spes Medica, Genova, Italy) was used, whereas for the rest of the participants an in-house developed strain gauge breathing sensor (Centre for Interdisciplinary Brain Research, University of Jyväskylä, Finland) that enables more accurate monitoring of breath holding was used.

### Preprocessing

First, MEG data were processed with MaxFilter™ 3.0 software (MEGIN, Helsinki, Finland). The temporal extension of the signal-space separation (tSSS) method was applied to remove the external magnetic interference from the MEG data (Taulu, Kajola, & Simola, 2004; Taulu & Kajola, 2005; Taulu & Simola, 2006). Bad channels were automatically detected and reconstructed in the MaxFilter software. Furthermore, the head position was estimated using 200 ms time windows with 10 ms steps for head movement compensation and the median head position across all participants’ MEG recordings was used for head position transformation. MEG data was also downsampled with a factor of 2 resulting in the sampling rate of 500 Hz. After preprocessing the collected MEG data was exported to Meggie that is an MNE-python-based graphical user interface (Heinilä & Parviainen, 2022 [preprint]) where common artifacts, such as eye blinks, horizontal eye movements, and cardiac artifacts were extracted, visually inspected, and manually removed using independent component analysis (ICA). The phases of the respiratory cycle were defined from the respiratory signal using MNE-python findpeaks function in which minima represent the onsets of inspiration and maxima the onsets of expiration. All the annotations created with this function were visually inspected and corrected, when necessary (e.g., onset of inspiration or expiration missing), using the annotation tool in Meggie. Thereafter these annotations were transformed into events (i.e., onsets of inspiration and expiration) used in the further analyses. Mean respiratory rate (breaths/minute) was determined based on the number of detected respiratory cycles during each breathing condition. Due to poor signal quality, we were not able to define all the phases of square breathing condition accurately and thus, this breathing condition was not included in the analysis concerning the phases of respiration. All further analyses were conducted using the gradiometers.

### Breathing techniques: Dynamic imaging of coherent sources and Fitting oscillations and one over f

To investigate the spatial distribution of modulations in periodic and aperiodic neural activity during spontaneous and volitionally controlled breathing, cross spectral density (CSD) matrices were first calculated for the frequency range of 2 – 45 Hz (length of the window 2048, 50% overlap). Next, dynamic imaging of coherent sources (DICS; Gross, Kujala, Hämäläinen, Timmermann, Schnitzler, & Salmelin, 2001) was applied to estimate the distribution of oscillatory activity. Since the DICS analysis was conducted at the parcel-level, the forward model of one participant (a single-compartment realistic boundary element model, fsaverage-5.1.0 template brain) was used to define the distribution of points across the parcels for all participants. As the parcellation we used a customized version of the Destrieux anatomical parcellation (Destrieux, Fischl, Dale, & Halgren, 2010) with 69 parcels per hemisphere that was constructed using a merge-and-split approach to produce uniform-sized parcels (Ala-Salomäki, Kujala, Liljestörm, & Salmelin, 2021). Thereafter, frequency-domain beamformer was implemented using common weights approach, in which the weights were defined by combining the CSD matrices of all the breathing conditions. Beamformer estimates at the parcel-level across the whole brain were calculated for all participants in each breathing condition (spontaneous, deep, and square breathing). Finally, Fitting oscillations and one over f (FOOOF) algorithm (Donoghue et al., 2020) was used to extract both periodic (i.e., power) and aperiodic components (i.e., exponent and offset) of the power spectra in each parcel within the frequency range of 2 – 45 Hz. For periodic activity, the power within alpha frequency band (8 – 13 Hz) was defined from the flattened spectra that was calculated by subtracting the aperiodic activity from the original spectra.

### Phases of breathing: Event-related dynamic imaging of coherent sources

Event-related dynamic imaging of coherent sources (erDICS; Laaksonen, Kujala, & Salmelin, 2008) was applied to explore the spatial distribution of suppressions and increases of oscillatory activity during inspiration and expiration. First, the time-frequency windows of interest for the source-level computations were defined. The frequency range was restricted to alpha frequency band (8 – 13 Hz) and to investigate the transitions from inspiration to expiration and vice versa, the time range of 0 – 2000 ms with a baseline of −500 ms – 0 ms with respect to the onset of inspiration and expiration was selected. Second, to move from sensor-level to source-level, CSD matrices were computed for a frequency range of 4 – 30 Hz and for a time window of −1000 ms – 2000 ms with respect to the onset of inspiration and expiration using a sliding window of 500 ms. Next, the CSD matrices were combined with the geometry of the participant’s head (a single-compartment realistic boundary element model, fsaverage-5.1.0 template brain) and the power distribution was estimated throughout the brain. Source reconstruction using a frequency-domain beamformer was implemented using common weights approach, in which the weights were defined by combining the CSD matrices of both inspiration and expiration within the same time intervals. Beamformer estimates across the whole brain were calculated for all participants in each breathing condition (spontaneous and deep breathing) and in selected time-frequency windows of interest (8 – 13 Hz, −500 ms – 2000 ms with respect to the onsets of inspiration and expiration).

### Statistical analyses

Statistical analyses of respiratory measures were performed using IBM SPSS Statistics 28.0 (IBM Corp., Armonk, New York, USA). First, descriptive statistics (mean, standard deviation) were calculated. Second, the normality of each variable was visually inspected using histograms and statistically tested with Kolmogorov-Smirnov test. Since the respiratory measures were not normally distributed, non-parametric Friedman test was performed to investigate the differences in mean respiratory rate between breathing conditions. Post hoc analyses were conducted using Wilcoxon Signed-Rank test with a Bonferroni correction resulting in a significance level of p < 0.017.

To compare periodic and aperiodic activity between different breathing techniques on the parcel-level, dependent samples t-test statistics were calculated using MATLAB (R2020b, The MathWorks Inc., Natick, MA, USA). Measures of periodic (alpha power) and aperiodic activity (exponent and offset) were compared between breathing techniques (spontaneous vs. deep, spontaneous vs. square, deep vs. square). The p-values (p < 0.01) across parcels were corrected for multiple comparisons using false discovery rate (FDR). Finally, the statistically significant results were visualized using FreeSurfer.

To estimate the statistically significant differences between the baseline and phases of breathing in different breathing conditions, a source-level cluster-based permutation test with a maximum-statistics approach was applied using MATLAB (R2020b, The MathWorks Inc., Natick, MA, USA). First, dependent samples t-test statistics comparing the distributions of power levels in baseline and inspiration/expiration time segments were calculated in each voxel. Voxels in which t-statistics exceeded an uncorrected threshold of α = 0.01 were clustered and new test statistics for each cluster were calculated by summing the t-statistics of the voxels included in the same cluster. Second, the samples were permuted randomly by exchanging the labels between the baseline and inspiration/expiration in a random number of participants. The total number of random permutations was 10 000 (5000 permutations for positive and 5000 permutations for negative values). Maximum-statistics approach with a corrected threshold of α = 0.01 was applied to reduce the amount of type I errors (i.e., false positives) due to multiple comparisons. Thus, maximum and minimum t-values of the test distributions in each cluster were collected and the final p-values were estimated by comparing the cluster-level t-values to the maximum/minimum distributions. This approach allowed us to distinguish between relative decreases and increases in the alpha band oscillatory activity during inspiration and expiration in comparison with the baseline. Finally, the statistically significant cluster-based permutation test results were visualized using FreeSurfer.

## Data availability

The data cannot be made openly available according to the ethical permission and national privacy regulations at the time of the study, but the data supporting the findings of this study are available upon reasonable request from the authors and with permission of the Ethics Committee of the University of Jyväskylä.

## Code Availability

The scripts used to perform the analyses are available upon request from the authors.

